# The Cell Surface Receptors Ror1/2 Control Cardiac Myofibroblast Differentiation

**DOI:** 10.1101/2021.03.02.433549

**Authors:** Nicholas W. Chavkin, Soichi Sano, Ying Wang, Kosei Oshima, Hayato Ogawa, Keita Horitani, Miho Sano, Susan MacLauchlan, Anders Nelson, Karishma Setia, Tanvi Vippa, Yosuke Watanabe, Jeffrey J. Saucerman, Karen K. Hirschi, Noyan Gokce, Kenneth Walsh

**Author notes:** Authors contributed equally to this work (Nicholas W. Chavkin, Soichi Sano). Corresponding Author: Kenneth Walsh, Ph.D., Address: University of Virginia School of Medicine, MR5 Building Room 1312, 415 Lane Road, Charlottesville, Virginia 22908.

## Abstract

**Background:** A hallmark of heart failure is cardiac fibrosis, which results from the injury-induced differentiation response of resident fibroblasts to myofibroblasts that deposit extracellular matrix. During myofibroblast differentiation, fibroblasts progress through polarization stages of early pro-inflammation, intermediate proliferation, and late maturation, but the regulators of this progression are poorly understood. Planar cell polarity receptors, receptor tyrosine kinase like orphan receptor 1 and 2 (Ror1/2), can function to promote cell differentiation and transformation. In this study, we investigated the role of the Ror1/2 in a model of heart failure with emphasis on myofibroblast differentiation.

**Methods and Results:** The role of Ror1/2 during cardiac myofibroblast differentiation was studied in cell culture models of primary murine cardiac fibroblast activation and in knockout mouse models that underwent transverse aortic constriction (TAC) surgery to induce cardiac injury by pressure overload. Expression of Ror1 and Ror2 were robustly and exclusively induced in fibroblasts in hearts after TAC surgery, and both were rapidly upregulated after early activation of primary murine cardiac fibroblasts in culture. Cultured fibroblasts isolated from Ror1/2-KO mice displayed a pro-inflammatory phenotype indicative of impaired myofibroblast differentiation. Although the combined ablation of Ror1/2 in mice did not result in a detectable baseline phenotype, TAC surgery led to the death of all mice by day 6 that was associated with myocardial hyper-inflammation and vascular leakage.

**Conclusions:** Together, these results show that Ror1/2 are essential for the progression of myofibroblast differentiation and for the adaptive remodeling of the heart in response to pressure overload.

## Introduction

Excessive fibrosis during pathological cardiac remodeling is a hallmark of heart failure and the response to cardiac injury. Cardiac fibrosis is the result of resident cardiac fibroblasts that undergo myofibroblast differentiation to promote inflammation and secrete extracellular matrix in response to various forms of cardiac injury^1–4^. Although injury-induced inflammation and matrix deposition is an adaptation to acute cardiac injury that can prevent the heart from rupture^5, 6^, excessive fibrotic deposition leads to ventricle stiffness that impairs cardiac function^7–9^. Recently, myofibroblast differentiation has been characterized as a transition through what is referred to as three phenotypic polarization stages: an initial pro-inflammatory phenotype, an intermediate proliferative phenotype, and a final mature phenotype^10^. These polarization stages have specific functional differences. The pro-inflammatory fibroblast recruits inflammatory cells to the cardiac tissue in response to injury, the proliferative fibroblast undergoes cell division and deposits extracellular matrix in response to TGF-β, and the mature myofibroblast maintains and strengthens the fibrotic deposits by expressing matrix remodeling proteins^11^. A recent study using single cell RNA sequencing to analyze murine interstitial cells after myocardial infarction revealed fibroblast populations consistent with these polarization states^12^. Despite its importance, we have limited knowledge of the factors that control the passage of myofibroblast through the different stages of differentiation.

The Receptor Tyrosine Kinase Like Orphan Receptors 1 and 2 (Ror1 and Ror2) have been evaluated for their roles in development and cell transformation^13–15^. These membrane proteins are most highly expressed in developing tissues^16^, and mice lacking Ror1 display skeletal defects^17^ whereas mice lacking Ror2 have impaired heart and limb development^18^. Ror1 and Ror2 expression can be reactivated in various cancers^19–24^, and knockdown or Ror1, Ror2, or both Ror1/2 in various transformed cell lines reduces their proliferation and migration^25–28^. Additionally, Ror1 expression in satellite cells promotes proliferation and skeletal muscle regeneration after injury^29^. These functions of Ror1/2 may be linked to the ability of this receptor system to control the planar cell polarity signaling pathway. Planar cell polarity is the asymmetrical alignment of cells to coordinate directionality with neighboring cells and extracellular matrix^30^. Proteins in the planar cell polarity pathway, including Ror1/2 and others (Ptk7, Prickle1, Vangl1, Vangl2, Dvl1, Dvl2, and Dvl3), regulate cell polarity by organizing the actin cytoskeleton and segregating proteins to opposite sides of the cell^31, 32^. In specific cell types, planar cell polarity can regulate proliferation, migration, and cell differentiation^33–38^. Thus it is tempting to speculate that planar cell polarity-mediated actin organization may be critical to myofibroblast function, and that it also functions as an integral step in the actin alignment-mediated signal transduction that promotes locomotion, contraction, and matrix reorganization during fibroblast differentiation^39^.

Although many of the functions attributed to Ror1 and Ror2 are shared by the process of fibroblast activation and differentiation, the roles of Ror1 and Ror2 in myofibroblasts has not been investigated previously. In the course of our studies, we found that Ror1/2 were generally expressed at low levels in the nonchallenged adult mouse tissues. However, the expression of these proteins become markedly in activated fibroblast in response to injurious stimuli. Thus, in this study, we investigated the role of Ror1 and Ror2 in cardiac remodeling through *in vivo* mouse models of heart failure and *in vitro* cell culture models of cardiac myofibroblast differentiation.

## Methods

### Mouse strains

All animal experiments were approved by the Animal Care and Use Committees at Boston University and the University of Virginia. Ror1/2^fl/fl^ + Ubc-Cre^ERT2^ mice were generated by combining Ror1 flox (Jackson Labs #018353,^40^), Ror2 flox (Jackson Labs #018354,^40^), and Ubc-Cre^ERT2^ (Jackson Labs #008085,^41^) alleles, which are all in the B6 mouse background. Additionally, Ror2 expression LacZ reporter mice (Ror2-LacZ,^42^) were used. Three mice per condition were used for fibroblast isolation experiments and five mice per condition were used for transverse aortic constriction experiments.

### Transverse Aortic Constriction (TAC) surgery

Cardiac pressure overload by transverse aortic constriction (TAC) was performed as previously described^43^. Briefly, surgery was performed on anesthetized mice where the aortic arch was accessed and constricted between the brachiocephalic artery and left common carotid artery. A 27-gauge spacer was placed parallel to the transverse aorta, and 8-0 vicryl suture (ETHICON Cat #J401G) was used to ligate the aorta. The spacer was removed, and the surgical wounds were sutured and mouse was allowed to recover. Successful TAC surgery was confirmed by initial expansion of the brachiocephalic artery and subsequent echocardiography measurements (FUJIFILM VisualSonics, Vevo 2100) at 0, 7, 14, and 28 days post-TAC. Fractional shortening was quantified by Cardiac sectioning was performed by cardiac tissue isolation, fixation with 10% neutral buffered formalin for 24 hours, tissue dehydration, then paraffin-embedding and sectioning. Cardiac tissue sections were either stained for 5-bromo-4-chloro-3-indolyl-b-galactosidase (Xgal; Roche Cat #XGAL-RO) or immunostained with rabbit anti-mouse polyclonal Ror2 (provided by Dr. Yasuhiro Minami, Kobe University) with DAB visualization (Vector Laboratories, Cat #SK-4100), and counterstained with hematoxylin and eosin (Sigma Aldrich, Cat #HHS128-4L and #HT110180-2.5L, respectively). Protein lysates from TAC heart samples were isolated from cardiac tissue by 1% Triton-X supplemented with protease inhibitors and probed for rabbit anti-mouse polyclonal Ror1 (provided by Dr. Michael Greenberg,^40^) rabbit anti-mouse monoclonal Ror2 (Cell Signaling Technology Cat #88639), or β-actin (Cell Signaling Technology Cat #4970) by western blot. Cardiac myocytes and cardiac fibroblasts were isolation in a langendorff apparatus as previously described^44^. These primary cells were either used for gene expression analysis after RNA isolation and purification using primers listed in **Supplementary Table S1**, or analyzed by flow cytometry using the following fluorescently-conjugated antibodies: CD45 (BioLegend Cat #103126), mEF-SK4 (Miltenyi Biotech Cat #103-102-352), Ror2 (R&D Systems Cat #AF2064).

### Primary cardiac cell isolation and fibroblast passaging

Interstitial cells of cardiac tissue were isolated by digestion and cell-type separation, based on previously published methods^45^. Cardiac tissue, either healthy, sham-surgery, or TAC surgery, was isolated from euthanized mice, minced with scissors, and digested in digestion media (HBSS with 0.1% Trypsin and 100 U/mL Collagenase Type II) for 80 minutes. CD31 + Endothelial Cells and CD45+ Leukocytes were separated and purified by FACS, with BV421 Rat anti-Mouse CD31 antibody (BD Biosciences Cat #562939) and PerCP Rat anti-Mouse CD45 antibody (BD Biosciences Cat #561047). Fibroblasts were purified by plating on plastic tissue culture dishes for 2 hours in Fibroblast Growth Medium 3 (PromoCell Cat #c-23130), then media was refed to eliminate cells that had not stuck, leaving the fibroblasts on the tissue culture plate. Genes were assessed by RNA isolation (Qiagen RNeasy Kit Cat# 74104), reverse transcriptase conversion to DNA (Applied Biosystems cDNA Reverse Transcription Kit Cat #4368814), and SYBR Green detection (Applied Biosystems Fast SYBR Green Master Mix Cat #43-856-16) with a quantitative Polymerase Chain Reaction (qPCR) machine (Applied Biosystems QuantStudio 6). Primers for Q-PCR are listed in **Supplementary Table S1**. Fibroblasts were further passaged and cultured in Fibroblast Growth Medium 3.

### Myofibroblast induction

Isolated cardiac fibroblasts at Passage (P)2 were treated with 10 ng/mL TGF-β1 (R&D Systems Cat #240-B) in Fibroblast Growth Medium 3 for four days to assess myofibroblast differentiation. Gene expression was assessed as described above using primers listed in **Supplementary Table S1**. Immunofluorescent staining of Acta2 (Cell Signaling Technologies Cat #36110) and DAPI (ThermoFisher Cat #62248) was performed by culturing cells in 4-well Chamber slides (ThermoFisher Cat #154526PK) with 10 ng/mL TGF-β1 in Fibroblast Growth Medium 3 for four days, fixation with 4% PFA for 20 min, permeabilization with PBS-T (PBS + 0.1% Tween20), incubation with fluorescent Acta2 antibody, then counterstain with DAPI. Immunostained cells were imaged by confocal microscopy (Leica SP8). Acta2 fiber alignment was quantified by ImageJ (Ver 2.0.0).

### Bulk RNA sequencing

Primary cardiac fibroblasts were isolated as described above from either control mice (Ror1/2^fl/fl^) or Ror1/2-KO mice (Ror1/2^fl/fl^ + Ubc-CreER^T2^) after injection with tamoxifen (Sigma-Aldrich Cat# T5648) when mice were 6-to 8-weeks old. Each isolation pooled four hearts from mice of the same litter, with the same male:female ratio variation between Control and Ror1/2-KO samples. Mice were bred between a Ror1/2^fl/fl^ genotype and a Ror1/2^fl/fl^ + Ubc-CreER^T2^ genotype to generate ~50% Control mice and ~50% Ror1/2-KO mice per litter, allowing for littermate paired samples. Specifically, isolations of Control and Ror1/2-KO mice were performed on four different litters with paired samples from the same litters (Ex: Control Sample 1 and Ror1/2-KO Sample 1 used hearts from mice of the same litter). Primary cardiac fibroblasts were grown for 9 days in culture, then RNA lysate was isolated and purified. Purified RNA was submitted to the University of Virginia Genome Analysis and Technology Core for whole transcriptome sequencing by first library preparation with NEBNext Ultra II Directional RNA Library Prep Kit for Illumina (NEB Cat #E7760) and then sequencing by Illumina NextSeq 500 Sequencing System for paired-end 75bp reads. Programs were used for bioinformatic analysis: raw read data was quality checked with FastQC (Babraham Bioinformatics), aligned with Kallisto^46^, analyzed for gene expression and differential expression with Sleuth^47^, and analyzed for GO term enrichment with GAGE^48^. Aligned and normalized sequencing results and GO term enrichment results are provided in **Supplementary Data 1-2**.

### Immunophenotyping of cardiac tissue

Cardiac tissue was isolated after Sham or TAC surgery. Tissue was digested with collagenase I (450 U/ml), collagenase XI (125 U/ml), DNase I (60 U/ml), and hyaluronidase (60 U/ml), and the isolated cells from digested tissue were immunostained with fluorescently-conjugated antibodies against CD45-Pacific Blue, 30-F11 (BioLegend Cat #103126), Ly6G-PE, 1A8 (BioLegend Cat #127618), CD11b-APC-Cy7, M1/70 (BioLegend Cat #101226), F4/80-PE-Cy7, BM8 (BioLegend Cat #123114), and Ly6C-FITC, HK1.4 (BioLegend Cat #128006). Dead cells were excluded by staining with DAPI. Immunostained fluorescent cells were analyzed by Flow Cytometry using BD LSR II Flow Cytometer (BD Bioscience). Additionally, isolated cardiac tissue was lysed and analyzed for inflammatory cytokines Il1b, Il6, and Ccl2 by qRT-PCR. Realtime PCR primers are listed in **Supplementary Table S1**.

### Evans Blue staining

Vascular permeability of the heart after pressure-overload was evaluated by the extent of the leakage of Evans blue dye (Sigma Aldrich Cat# E2129). Mice were sacrificed 30 minutes after tail-vein injection of 1% w/v in 0.9% saline. Dyes were allowed to circulate throughout the body during this period.

### Statistical analysis

Unless otherwise indicated, statistical analysis was performed using either a standard two-tail Student’s t-test or a two-way ANOVA test followed by a Tukey’s multiple comparison corrected post-hoc test by GraphPad Prism (GraphPad Software, Inc., San Diego, CA). All statistical analysis of RNA sequencing datasets was performed through computational analysis packages, which contain statistical corrections for large data sets.

## Results

### Transverse aortic constriction induces early Ror1/2 expression in cardiac tissue

Cardiac pressure overload in the mouse model of transverse aortic constriction (TAC) induces fibroblast activation and myocardial remodeling, including initial inflammation (1-3 days post-TAC), extracellular matrix remodeling (3-14 days post-TAC), and eventual heart failure (14-28 days post-TAC)^49^. Thus, cardiac tissue was analyzed before and after TAC surgery to investigate the expression patterns of Ror1 and Ror2 during the remodeling time course. The TAC surgery model was initially performed on Ror2-LacZ mice that has the *LacZ* reporter gene knocked-in to the *Ror2* locus^42^. There was no detectable Ror2-mediated *LacZ* expression in uninjured, or sham-operated mice (**Figure 1A** and not shown), but β-galactosidase expression was markedly induced at 7 days post-TAC (**Figure 1B**). Immunohistochemical analysis revealed that Ror2 protein was concentrated in cells of the interstitial space of the myocardial tissue at 7 days post-TAC (**Figure 1C**). Next, we determined the time course of Ror1 and Ror2 protein induction after TAC surgery by western blot analysis of protein lysates from cardiac tissue. The expression of both Ror1 and Ror2 protein was increased by three-days post-TAC, peaking at 7-days, and then decreasing at 14- and 28-days post-TAC (**Figure 1D**).

**Figure 1.**
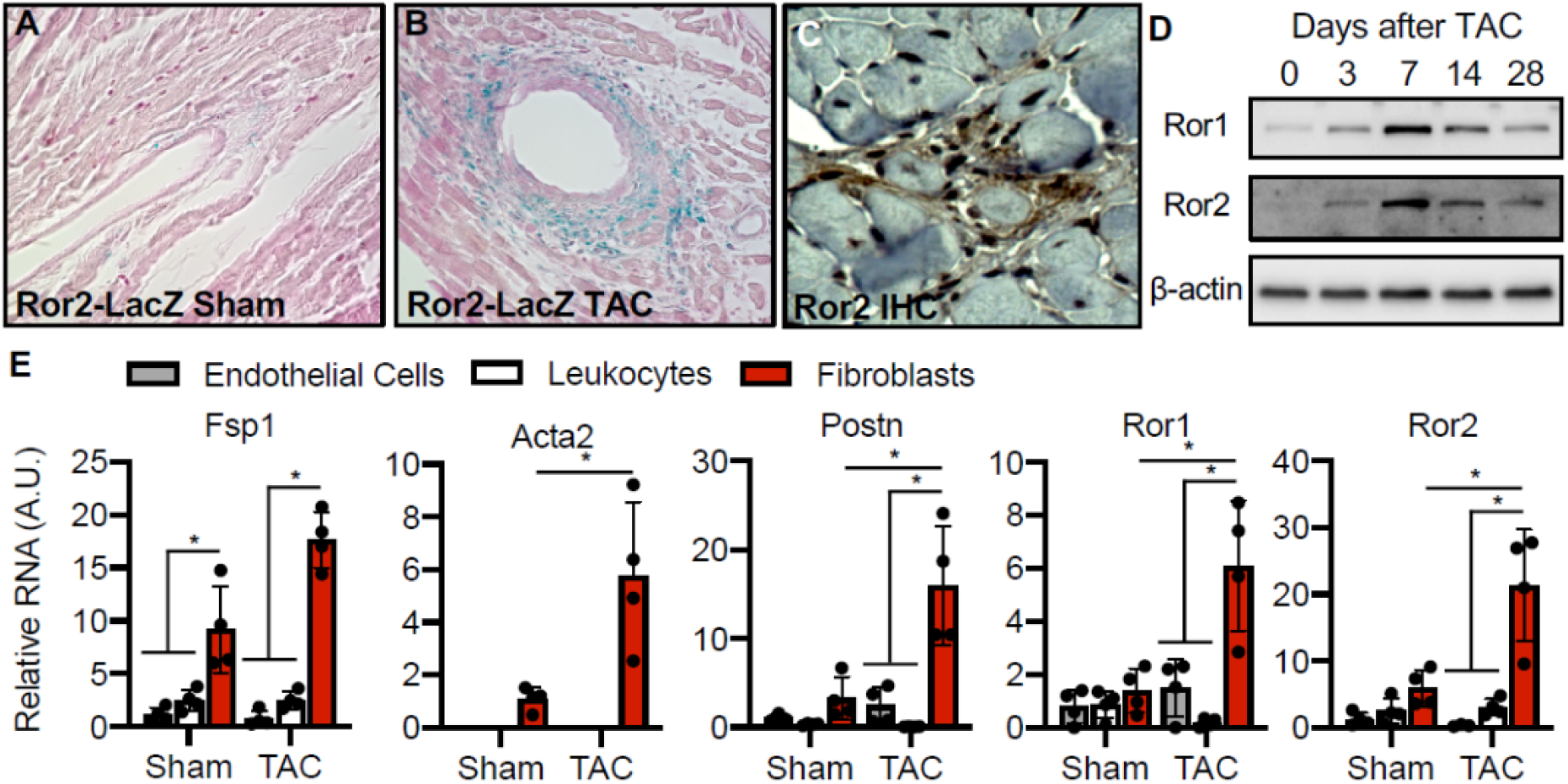
Ror1 and Ror2 expression after Transverse Aortic Constriction in mice. Cells expressing Ror2 in cardiac tissue was visualized in Ror2-LacZ mice 28 days after A) sham surgery or B) TAC surgery. C) Protein localization was visualized in cardiac tissue 7 days after TAC by IHC for Ror2. D) Ror1 and Ror2 protein expression was measured by western blot at various days after TAC surgery (β-actin as a loading control). E) Endothelial cells, leukocytes, and fibroblasts were isolated 7 days after Sham or TAC surgery, and RNA expression of relevant genes was quantified.

To determine the cell type(s) that express Ror1 and Ror2, RNA was isolated from endothelial cells, leukocytes, and fibroblasts from sham-operated and TAC-treated hearts 7 days post-surgery, and qPCR was performed to detect the levels of relevant transcripts. Ror1 and Ror2 transcript expression was detected in the activated cardiac fibroblasts at 7 days after TAC, but not in the endothelial or leukocyte cell populations (**Figure 1E, Supplemental Figure 1).** Ror2 protein expression was also detected in fibroblasts after TAC surgery by flow cytometry (**Supplemental Figure 2**). Overall, the timing, location, and cell-specific gene expression patterns suggest that both Ror1 and Ror2 are induced during the activation and expansion of cardiac fibroblasts that occurs in response to pressure overload-induced myocardial remodeling.

### Ror1/2 are induced during early cardiac fibroblast activation

The induction of Ror1/2 during myofibroblast differentiation was also investigated in cultured cells using isolated murine cardiac fibroblasts. Fibroblasts from wild-type murine hearts were attached to cell culture plates, flattened, and expanded over nine days post-isolation (**Supplemental Figure 3**). RNA expression of key genes increased at different rates during this activation time course (**Figure 2A**). As expected, Fsp1 was induced by day 3 and the fibroblast activation genes Slug and Snail increased at day 9. Both Ror1 and Ror2 increased over this time course, but the induction of Ror2 preceded that of Ror1. The planar cell polarity protein transcript Ptk7 increased by day 3 and continued to increase at days 6 and 9, but Prickle and Vangl2 displayed no statistically significant change in expression.

**Figure 2.**
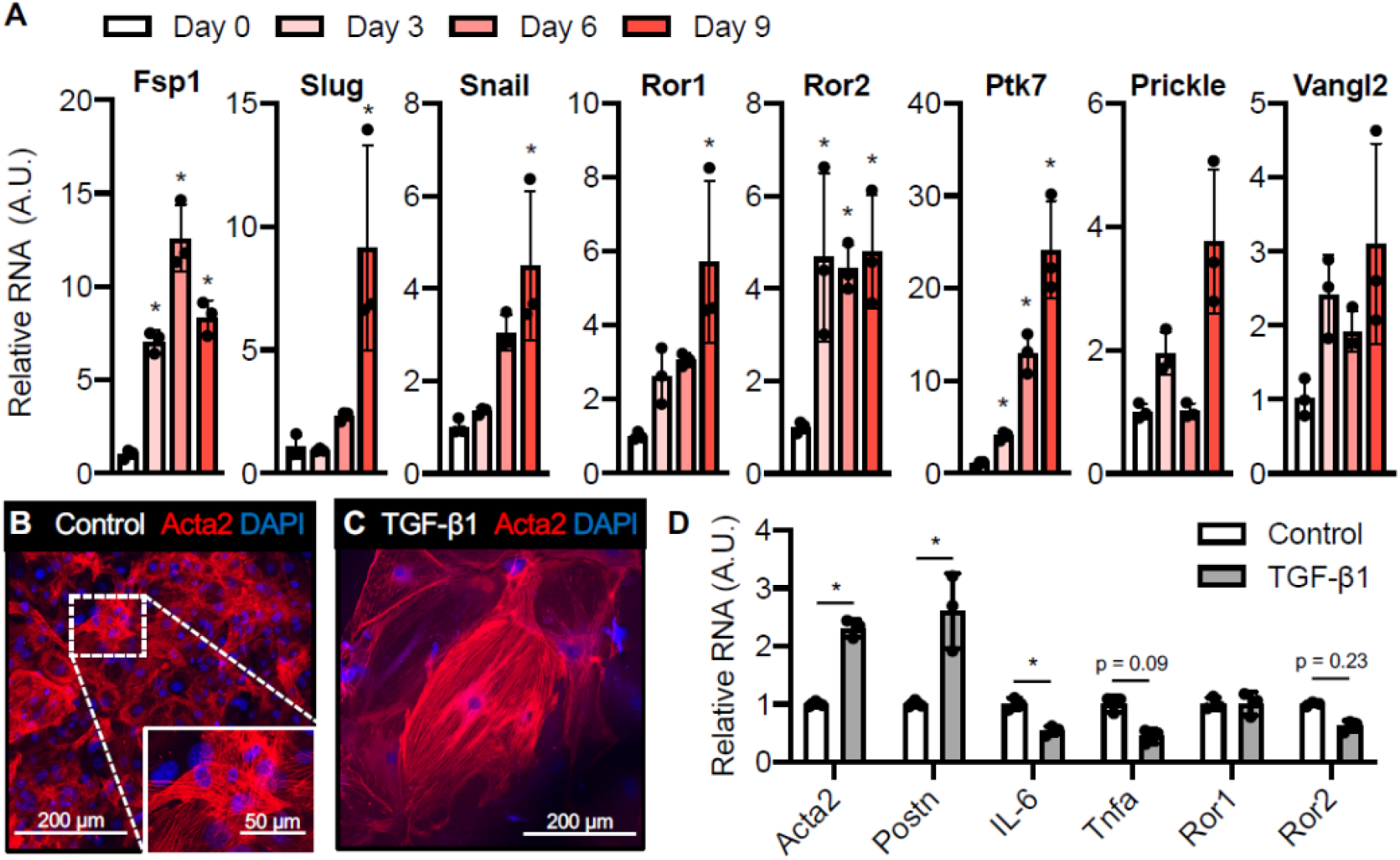
Induction of Ror1 and Ror2 during early fibroblast activation and induction of myofibroblast differentiation of murine cardiac fibroblasts. Murine cardiac fibroblasts were isolated from healthy cardiac tissue of C57Bl6 mice. A) RNA expression of genes associated with fibroblast identity, fibroblast activation, Ror1/2 receptors, and planar cell polarity signaling was quantified over time. Cardiac myofibroblasts at passage 2 were stained for SMØ-actin fibers after B) control and C) TGF-β1 treatment. D) RNA expression of genes associated with myofibroblast differentiation, inflammation, and Ror1/2 receptors was quantified in control and TGF-β1 treated fibroblasts.

Next, myofibroblast differentiation was stimulated in these cultures by treatment with TGF-β1. Myofibroblast differentiation could be detected by cell enlargement and SMα-actin filament elongation (**Figure 2B-C**). However, in contrast to the earlier stage of fibroblast activation, TGF-β1 treatment did not lead to a further increase in Ror1 or Ror2 expression. On the other hand, hallmarks of myofibroblast differentiation could be observed following TGF-β1 treatment including the increased transcript expression of myofibroblast differentiation proteins Acta2 and Postn, and decreased expression of the cytokines IL-6 and Tnfα (**Figure 2D**). Together, these data suggest that the induction of Ror1/2 is an early event during fibroblast activation, and that their expression is not altered at the later stages of myofibroblast differentiation.

### Ror1/2 double-knockout fibroblasts exhibited an immature cardiac myofibroblast phenotype

We next investigated the role of Ror1/2 in fibroblast activation in the fibroblast cell culture system using primary cardiac fibroblast cells isolated from control or Ror1/2-KO mice. Ror1/2 was eliminated in adult mice through an inducible transgenic knockout strategy by crossing Ror1/2^fl/fl^ and Ubc-CreER^T2^ mice and treating the progeny with tamoxifen for 2 weeks. These mice displayed no observable phenotype, and they had a normal cardiac phenotype three months after the induction of gene ablation (**Supplemental Figure 4**). Primary cardiac fibroblasts were isolated from control or Ror1/2-KO mice and grown to confluence. RNA was then isolated, and next-generation RNA sequencing was performed. Many genes associated with a manually curated network of fibrosis and fibroblasts^50, 51^ were differentially regulated between control and Ror1/2-KO fibroblasts (**Figure 3A**). As expected, cells were largely void of Ror1 and Ror2 transcripts, and the planar cell polarity gene Ptk7, *Vangl1, Vangl2, Prickle1, Dvl1, Dvl2, Dvl3*) were generally reduced in the Ror1/2-KO fibroblasts. Notably, Ror1/2-KO fibroblasts showed an increase in inflammatory cytokine gene expression (*Il1a, Il6, Tnfa, Ccl2*) and a decrease in transcripts associated with cell proliferation (*Pcna, Mki67, Cdk1, Cdk2, Cdk4, Cdk6*). Additionally, *transcripts associated with the promotion of* fibrosis in the ERK1/2 signaling and matrix remodeling pathways were downregulated (*Map2k3, Map3k1, Map3k5, Mapk14, Mapk8, Timp1, Timp2*), and transcripts associated with the inhibition of fibrosis in the matrix remodeling and natriuretic signaling pathways were upregulated (*Mmp2, Mmp9, Mmp14, Nppa, Nppb, Npr1, Npr2, Npr3*). Analysis of the top 10 upregulated and downregulated Gene Ontology terms between control and Ror1/2-KO fibroblasts revealed the strong induction of inflammation-related pathways and an inhibition of proliferation-related and microtubule organization-related pathways (**Figure 3B**). The specific genes included in these groupings were highly differentially expressed between control and Ror1/2-KO fibroblasts (**Figure 3C**). To the extent that the cell culture system models the early activation of fibroblasts, Ror1/2 dual deficiency led to the upregulation of inflammation pathways and down-regulation of pathways associated with matrix production, proliferation, and microtubule organization. Collectively, these data are consistent with a cellular fibroblast phenotype that appears to be stalled in the pro-inflammatory polarization state, suggesting that Ror1/2 are required for the progression of fibroblasts from this immature state to the intermediate proliferative state^10^.

**Figure 3.**
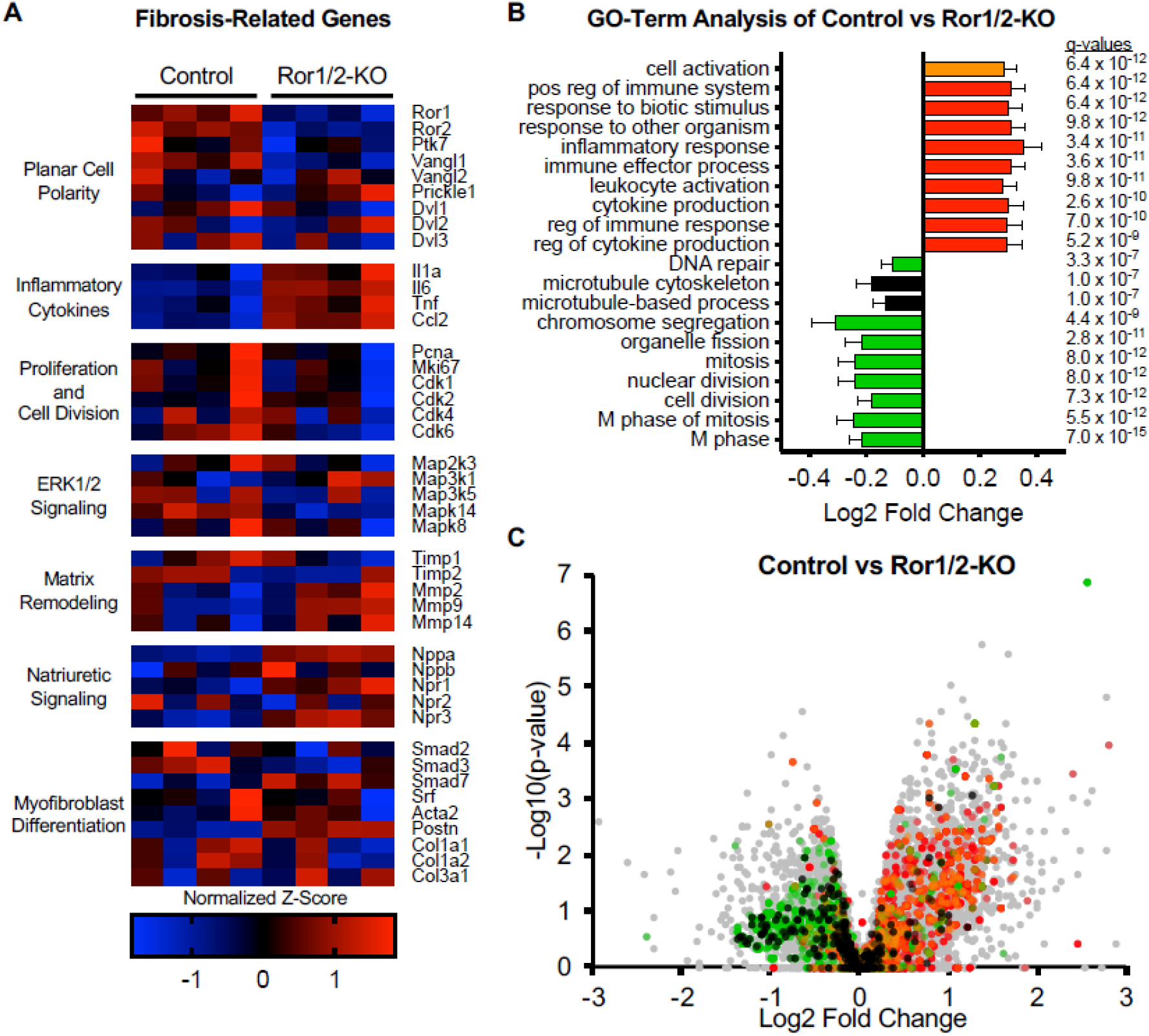
Ror1/2-mediated early fibroblast activation in primary murine cardiac fibroblasts. Primary cardiac fibroblasts were isolated from healthy control and transgenic Ror1/2 double-knockout mice transcriptional phenotype was assessed by bulk RNA sequencing. A) RNA transcript reads of specific genes related to planar cell polarity, inflammatory cytokines, proliferation and cell division, ERK1/2 signaling, matrix remodeling, natriuretic signaling, and myofibroblast differentiation were quantified as normalized Z-score (4 samples for each genotype). B) Gene ontology analysis of differential gene expression between control and Ror1/2-KO cardiac fibroblasts showed the top 10 up-regulated and down-regulated terms by q-value, with terms grouped by color: cell activation in orange, inflammation in red, proliferation in green, and microtubule regulation in black. C) Significance versus fold change of each gene between control and Ror1/2-KO samples was visualized by volcano plot, with genes in each gene ontology term highlighted in corresponding colors (all other genes in grey).

### Ror1/2 double-knockout fibroblasts are less responsive to TGF-β1 induced myofibroblast maturation

Previous findings showed that activated fibroblasts in the early pro-inflammatory state are less responsive to TGF-β stimulation than fibroblasts in the intermediate proliferative state^10^. Thus, to investigate whether the Ror1/2-KO fibroblasts are functionally stalled in an initial pro-inflammatory phenotype, we compared the responsiveness of control and Ror1/2-KO fibroblasts to TGF-β1-induced myofibroblast differentiation. In control fibroblasts, TGF-β1 treatment led to gene expression changes that are consistent with myofibroblast differentiation (increased Acta2 and Postn, decreased IL-6). However, Ror1/2-KO fibroblasts displayed diminished Acta2 and Postn induction and no decrease in IL-6 after TGF-β1 treatment when compared to control fibroblasts (**Figure 4A, Supplemental Figure 5**).

**Figure 4.**
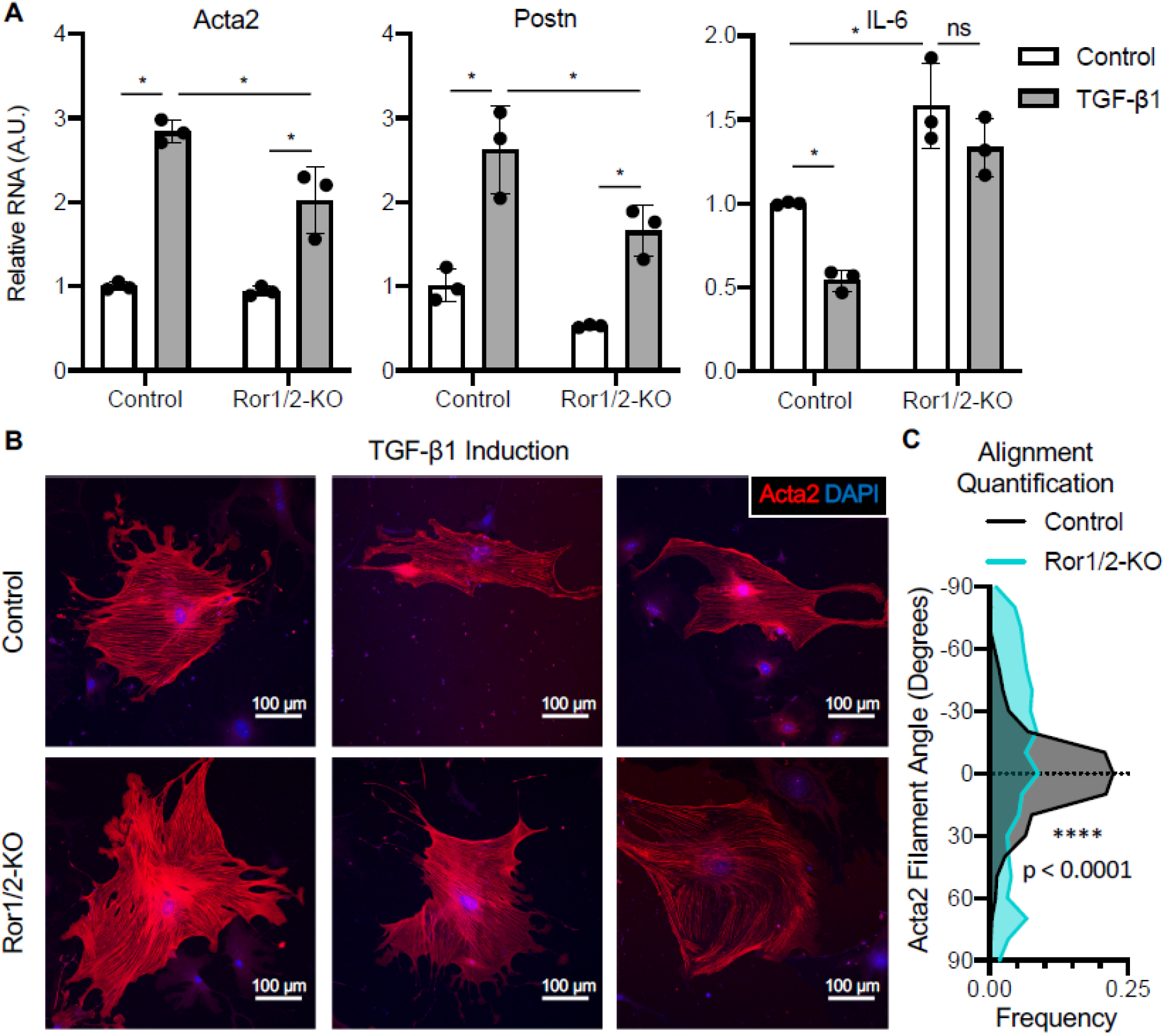
Ror1/2-mediated response to myofibroblast differentiation induced by TGF-β1. Myofibroblast differentiation was induced by treatment with 10ng/mL TGF-β1 for 4 days. A) RNA expression of myofibroblast-related (Acta2 and Postn) and inflammation-related (IL-6) genes was quantified in control and Ror1/2-KO fibroblasts. B) SMΓ]-actin filaments were visualized by immunofluorescent staining of sub-confluent cells in control and Ror1/2-KO fibroblasts, and C) alignment of SMΓ1-actin filaments was quantified and normalized per cell.

Next, TGF-β1-treated control and Ror1/2-KO fibroblasts were visualized after Acta2 immunostaining. Although treatment with TGF-β1 induced similar fibroblast cell enlargement between control and Ror1/2-KO fibroblasts (Control = 0.073±0.016 mm^2^, Ror1/2-KO = 0.071±0.010 mm^2^; p = 0.92), the Acta2 fibers appeared mis-aligned and disorganized in the Ror1/2-KO fibroblasts compared to the Acta2 fiber alignment in the control fibroblasts (**Figure 4B**). Quantification of the directional angle of individual Acta2 fibers within each cell showed that the Acta2 fibers in the Ror1/2-KO fibroblasts were significantly less aligned with each other than the Acta2 fibers in the control fibroblasts (**Figure 4C**). Taken together, these results show that Ror1/2-KO fibroblasts are less responsive to TGF-β1, a feature that is consistent with the notion that these cells are in the less mature, pro-inflammatory state. These results further suggest that the planar cell polarity signaling pathway may be critical for actin alignment during fibroblast maturation.

### TAC induces hyper-inflammation, rapid heart failure, and death in Ror1/2 double-knockout mice

To investigate the role of Ror1/2 in cardiac tissue remodeling *in vivo*, the inducible Ror1/2 double knockout mice (Ror1/2^fl/fl^ + Ubc-CreER^T2^) were treated with tamoxifen for 2 weeks and subjected to TAC. Ror1/2 double knockout mice displayed a rapid decline in heart function after TAC, as observed by echocardiography measurements at 3 days post-TAC (**Figure 5A,B**). This rapid development of systolic dysfunction was associated with death of the Ror1/2 double knockout mice between 4- and 6-days post-TAC (**Figure 5C**). Consistent with the heart failure phenotype, the TAC-treated, Ror1/2 double-knockout mice displayed marked increases in heart and lung weights (**Supplemental Figure 6**).

**Figure 5.**
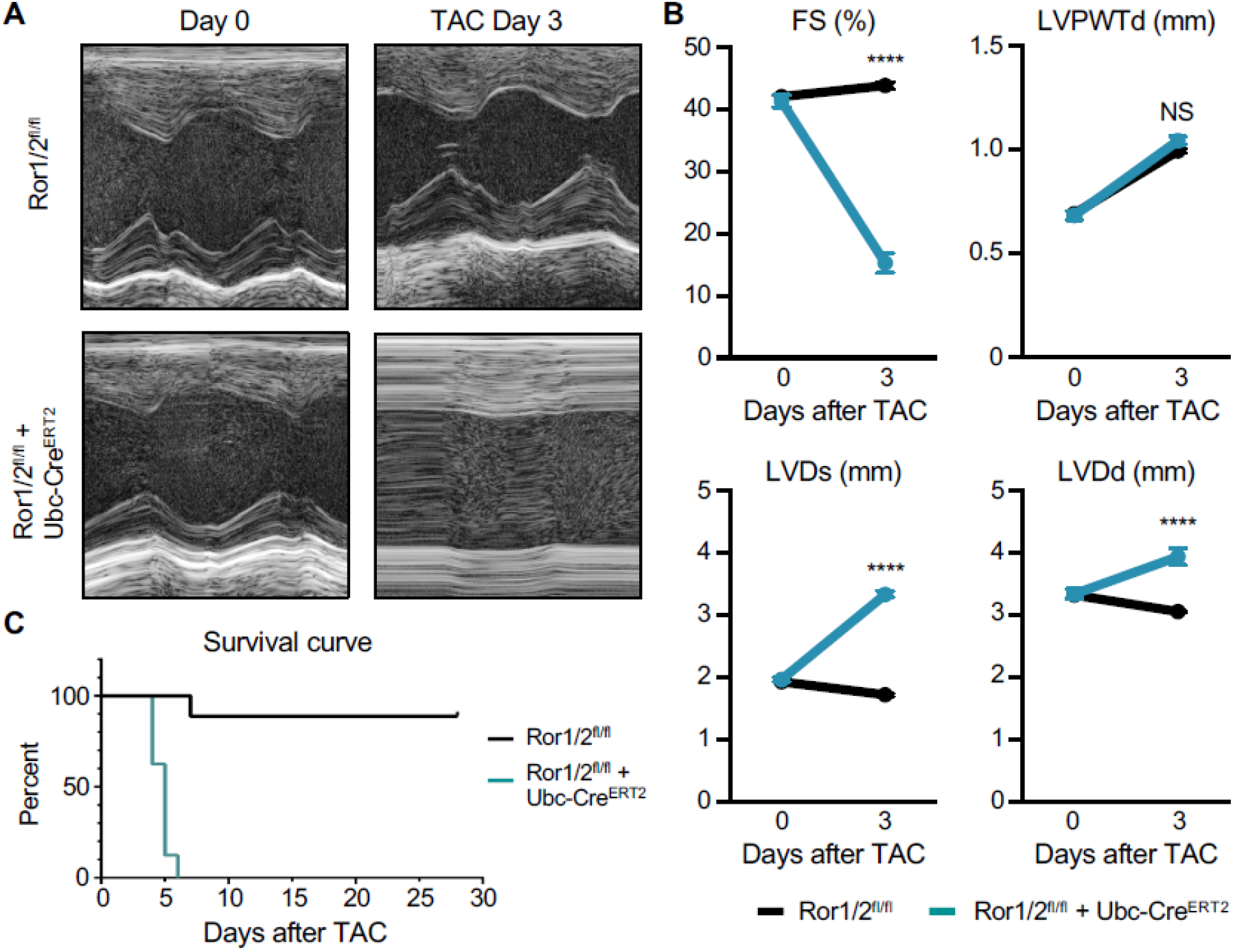
Early heart failure in transgenic Ror1/2 double knockout mice after TAC surgery. A) Transgenic Ror1/2 double knockout and control mice were subjected to TAC surgery, and cardiac output was imaged by echocardiography. B) Cardiac output factors were quantified: fractional shortening, left ventricular posterior wall thickness at end diastole, left ventricular diameter at end systole, and left ventricular diameter at end diastole. C) Survival of Ror1/2 double knockout mice and control mice after TAC surgery was recorded each day.

Hematoxylin and eosin staining of heart sections revealed a massive inflammatory infiltrate in the Ror1/2 double-knockout mice after TAC (**Figure 6A**). Inflammatory cell infiltration was further characterized by flow cytometry analysis of cells isolated from digested cardiac tissue at 3 days post-TAC. This analysis revealed a large increase in the number of CD11b+Ly6G+ neutrophils, CD11b+Ly6G-F4/80+ macrophages, and CD11b+Ly6G-Ly6C+ monocytes in the Ror1/2 double-knockout mice compared to wild-type mice treated by TAC (**Figure 6B-C**). In Ror1/2 double-knockout mice, TAC also caused a large induction of pro-inflammatory cytokine gene transcripts including *Il1b, Il6*, and *Ccl2* (**Figure 6D**). Notably, vascular leakage could also be detected after TAC in the Ror1/2 double-knockout mice as indicated by the permeability of the myocardium to Evans Blue dye (**Figure 6E**). These data reveal that the hearts of TAC-treated, Ror1/2 double-knockout mice are hyper-inflammatory. This phenotype is consistent with the notion that Ror1/2 is essential for the progression of fibroblast maturation from the initial pro-inflammatory stage during their activation sequence.

**Figure 6.**
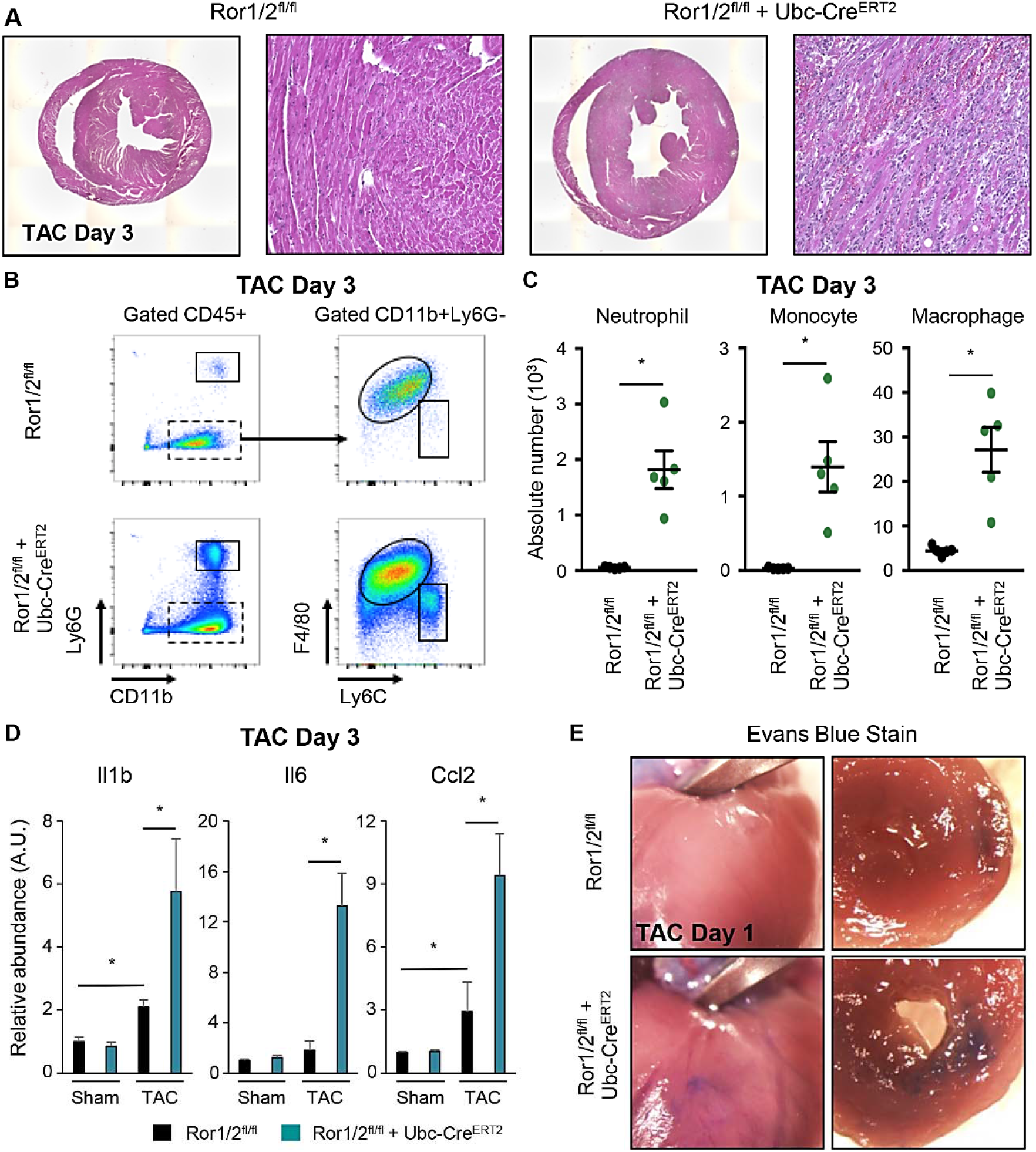
Inflammation response in transgenic Ror1/2 double knockout mice after TAC surgery. Inflammatory profile of control and transgenic Ror1/2 double knockout mice was assessed after TAC or sham surgery. A) H&E staining of cardiac tissues 3-days post-TAC were imaged. B) Cells were isolated from cardiac tissue and relative quantity of leukocyte populations were determined by flow cytometry, C) quantified by absolute number. D) Gene expression of pro-inflammatory cytokines in cardiac lysate was measured. E) Vascular permeability at 1-day post-TAC was assessed by Evans Blue dye injection, with vascular leakage visualized by blue dye in the cardiac tissue.

## Discussion

Through these experiments we examined the roles of the cell surface receptors Ror1 and Ror2 in cardiac myofibroblast differentiation. We found that Ror1/2 are upregulated in fibroblasts early after activation, both *in vivo* by pressure-overload cardiac injury induced by TAC surgery and in cardiac fibroblast cell culture. Multiple lines of data indicate that dual Ror1/2 deficiency impairs myofibroblast differentiation. Specifically, cultured Ror1/2-KO fibroblasts exhibited an immature, pro-inflammatory state and they were impaired in their response to TGF-β1 stimulation. Additionally, while mice lacking Ror1/2 did not exhibit a baseline phenotype, they were unable to acutely adapt to pressure overload cardiac injury. In response to TAC, these mice underwent profound decompensated remodeling by 3 days and they typically died within 6-days of surgery. Collectively, these results suggest that the early induction of Ror1/2 in fibroblasts is essential for the appropriate myofibroblast differentiation and required for the myocardium to adapt to the initial stages of pressure overload injury.

The phenotype of the Ror1/2 double-knockout mice highlighted a critical role for Ror1/2 in controlling the inflammatory phenotype of fibroblasts during the early stages of cardiac remodeling. The rapid upregulation of cytokines in response to TAC injury leads to the infiltration of neutrophils and monocyte-derived macrophages^43, 52^. Several recent studies have highlighted the role of fibroblasts in promoting inflammation during the initial phases of the cardiac remodeling response to myocardial infarction^10, 53^ and pressure overload^54, 55^. Our results expand on these studies by showing that fibroblasts lacking Ror1/2 appear to be phenotypically similar to fibroblasts in the transient, pro-inflammatory state that occurs early in the fibroblast differentiation continuum following experimental myocardial infarction^10^. Transcriptomic analyses of cultured cardiac fibroblasts revealed that Ror1/2-deficiency was associated with a decrease in pathways that promote fibrosis (proliferation genes, ERK1/2 signaling genes, tissue inhibitors of matrix metalloproteinases, and Smad2/3) and an increase in pathways that inhibit fibrosis (inflammatory cytokines, matrix metalloproteinases, and natriuretic signaling genes). These differences in gene expression suggest that the Ror1/2-KO fibroblasts are in an immature state that is pro-inflammatory, leukocyte-recruiting and less responsive to fibrotic stimuli. We confirmed this by showing that Ror1/2-KO fibroblasts were less responsive to TGF-β1 compared to control fibroblasts, displaying reduced induction of *Acta2* and *Postn* gene expression and diminished repression of *Il6* gene expression. Thus, we propose that Ror1/2-deficiency causes activated fibroblasts to stall in an early pro-inflammatory state of differentiation such that they are unable to efficiently transition to the intermediate proliferative state and mature homeostatic state. Also consistent with this hypothesis, we found that Ror1/2 double knockout mice exhibit a hyper-inflammatory phenotype associated with elevated cytokine transcript expression, exuberant inflammatory cell infiltration and vascular leakage within 3 days of TAC surgery. These results highlight the critical role of early Ror1/2 induction in the control of the early inflammatory response in the injured myocardium, and suggest that hyper-inflammatory activated fibroblasts can have a detrimental role in myocardial remodeling.

In myofibroblasts, appropriate SMα-actin stress fiber organization is required for the transduction of signaling responses to external stimuli^39^, and the misalignment of SMα-actin stress fibers is associated with dysregulated myofibroblast differentiation^56, 57^. Consistent with the notion that Ror1/2 induction is essential for myofibroblast differentiation, we find that Ror1/2-deficiency leads to SMα-actin filament misalignment in TGF-β1-treated fibroblasts. Ror1/2 signaling can control stress fiber alignment, myofibroblast differentiation and other cellular phenotypes through the regulation of upstream and downstream planar cell polarity components^13, 43, 58–63^. This system involves the participation of inner plasma membrane proteins associated with Ror1/2 that control cell polarity and asymmetric cell division through the regulation of actin filament organization by the action of the small GTPases RhoA and Rac^31, 32^. Our work is consistent with other studies that have implicated planar cell polarity proteins in myofibroblast differentiation, including the reported induction of Frizzled2 in myofibroblasts after experimental myocardial infarction^64^ and hypoxia-mediated suppression of myofibroblast differentiation through RhoA inhibition^65^. Furthermore, ablation of Smad3, a key signaling protein downstream of TGF-β cardiac fibroblasts^66^, will lead to the suppression of RhoA and a disruption of actin alignment^57^. Together, these results suggest a potential role of planar cell polarity signaling in mediating stress fiber organization and TGF-β response in activated cardiac fibroblasts.

We acknowledge that Ror1/2 double-deficient mice were constructed using a global knockout strategy that employed ubiquitin-CreER^T2^. As discussed, these mice lack a baseline phenotype, yet undergo rapid decompensated heart failure and death in response to pressure overload. Although we cannot rule out the contribution of other cell types lacking Ror1/2 to this phenotype, we note that Ror1/2 expression appears specific to activated cardiac fibroblasts under these conditions. Our experimental evidence in support of this include that expression of Ror1/2 was greatly increased after TAC-induced cardiac injury in the interstitial cells of the myocardium, and the TAC-induced upregulation of Ror1/2 gene expression was specific to fibroblasts with no Ror1/2 induction in endothelial cells or leukocytes. Consistent with these findings, a number of other studies have reported Ror1 and/or Ror2 expression in activated fibroblasts or fibroblast-like cells^12,67–70^, and a recent proteomic analysis of the human heart reported that ROR1 expression was 200-fold higher in than cardiac fibroblasts than other cardiac cell types^71^. In addition, studies with isolated fibroblasts further corroborate the in vivo observations. These *in vitro* studies documented the robust expression of Ror1/2 and associated planar cell polarity protein transcripts (Ptk7, Vangl1 and 2, Prickle1 and Dvl1, 2 and 3) in activated cardiac fibroblasts, and showed that the ablation of Ror1/2 leads to a proinflammatory phenotype that is consistent with the cardiac phenotype of the Ror1/2-deficient mouse following TAC surgery.

### Conclusion

Cardiac fibroblasts are sentinel cells in the heart that respond to early injury by adopting a proinflammatory and leukocyte-recruiting phenotype, followed by their transition to a reparative/proliferative phase that is pro-angiogenic and pro-fibrotic, and followed by a homeostatic phase. Our study reveals the critical role that the cell surface receptors Ror1/2 play in allowing myofibroblasts to appropriately transition through these phases. While the deficiency of these proteins has no detectable baseline phenotype, Ror1/2-deficient fibroblasts activated by pressure overload appears stalled in the early phase of the fibroblast differentiation continuum - a stage of differentiation that is highly pro-inflammatory. Further analysis of the injury-induced, hyper-inflammatory phenotype of the Ror1/2 double knockout mice may provide a greater window of understanding of how excessive inflammation contributes to pathological cardiac remodeling. Elevated inflammation is predictive of worse outcomes in patients with heart failure^72, 73^, and recent clinical trials of anti-inflammatory therapies targeting IL-1β and IL-1R1 have shown promising results in the treatment of this condition in some patient groups ^74–76^. Our results suggest that impairments in planar cell polarity-mediated regulation of myofibroblast differentiation could have a causative role in the development of excessive inflammation that is associated with heart failure. Therefore, a better understanding of the molecular mechanisms that regulate the progression of myofibroblast differentiation could provide opportunities for the development of therapies that more effectively reduce cardiac inflammation.

## Non-standard Abbreviations and Acronyms

Ror1/2: Receptor tyrosine kinase like orphan receptor 1 and 2
TAC: Transverse Aortic Constriction
TGF-β: Transforming Growth Factor Beta
Ptk7: Tyrosine-protein kinase-like 7 (Colon carcinoma kinase 4 (CCK4))
Prickle1: Prickle homologue 1
Vangl1/2: Vang-Like 1 and 2
Dvl1/2/3: Disheveled 1, 2, and 3
Ubc-Cre^ERT2^: Ubiquitin C-driven Cre recombinase with tamoxifen-inducible mutant human estrogen receptor (ERT2)
LacZ: Beta-galactosidase

## Acknowledgements

We would like to thank the University of Virginia Genome Analysis and Technology Core for advice and technical support on bulk RNA sequencing, and the University of Virginia Health Sciences Library and Research Computing Center for assistance with computational analysis.

## Supplemental Materials

Supplemental Figures 1-6

Supplemental Table 1

