## Supplemental Figures for "The Cell Surface Receptors Ror1/2 Control Cardiac Myofibroblast Differentiation"

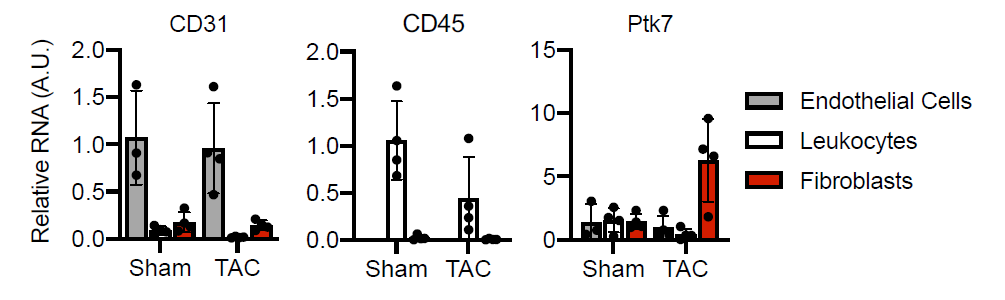

**Supplemental Figure 1.** Endothelial cells, leukocytes, and fibroblasts were isolated 7 days after Sham or TAC surgery, and RNA expression of relevant genes was quantified.

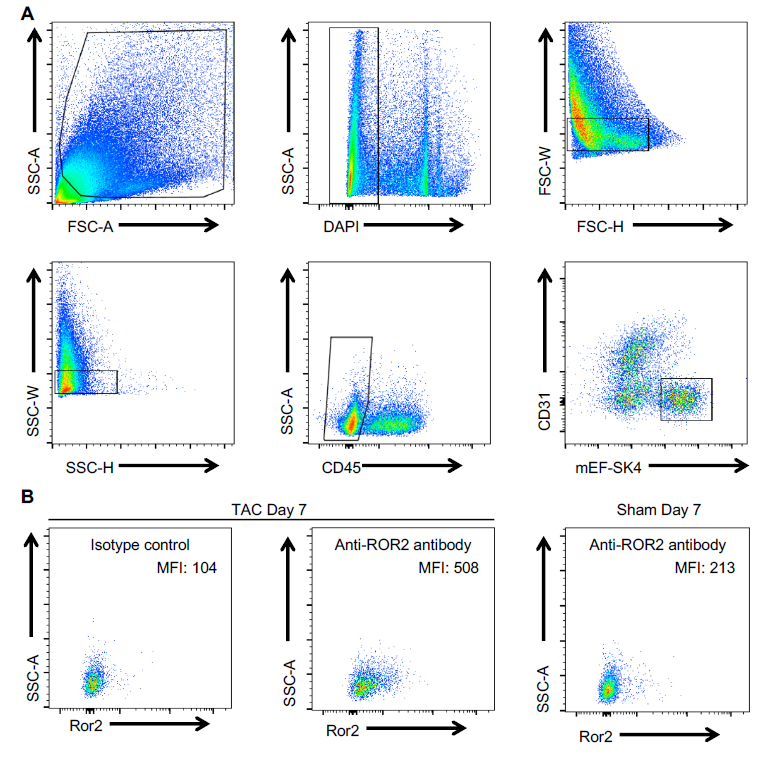

**Supplemental Figure 2.** A) Cardiac fibroblasts were isolated by langendorff apparatus and identified by flow cytometry using the gating strategy outlined. B) Ror2 expression in the fibroblast population was determined after TAC or Sham surgery (Antibody isotype control staining as a control), with Mean Florescence Intensity (MFI) reported.

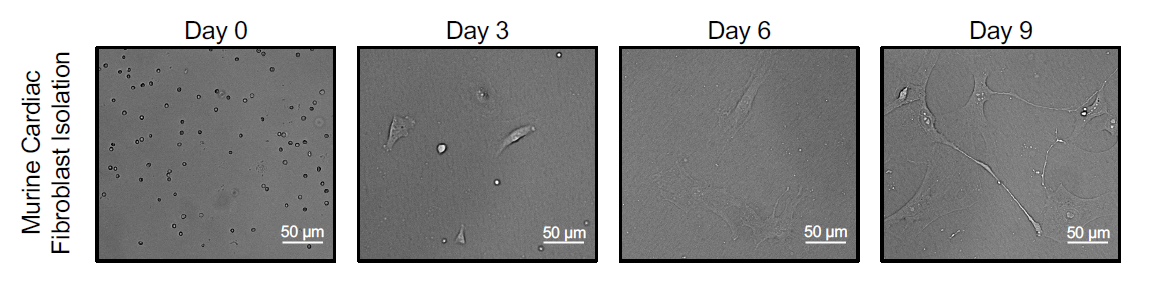

**Supplemental Figure 3.** Primary fibroblast cells from cardiac tissue were imaged at different time points while plated on tissue culture plastic to visualize early fibroblast activation.

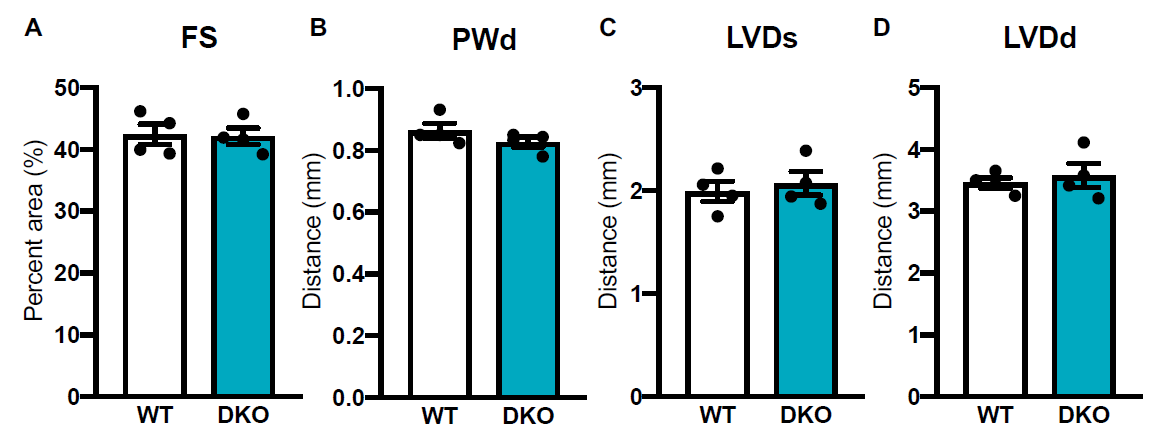

**Supplemental Figure 4.** Echocardiography measurements of functional cardiac parameters were quantified in Ror1/2^fl/fl^ (WT) and Ror1/2^fl/fl^ + Ubc-CreER^T2^ mice (DKO) 3 months after tamoxifen-induced DNA recombination: A) Fractional shortening, B) Posterior wall thickness at end diastole, C) Left ventricular diameter at end systole, and D) Left ventricular diameter at end diastole.

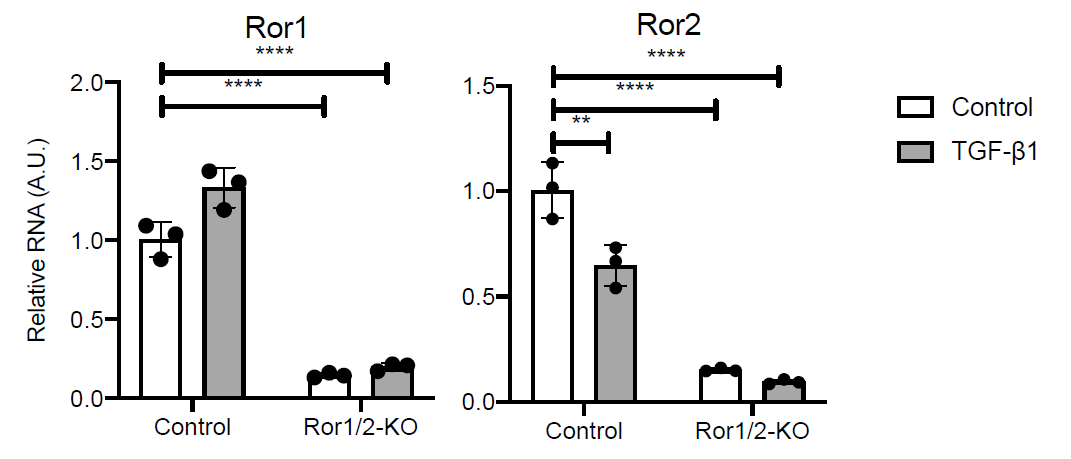

**Supplemental Figure 5.** RNA expression of Ror1 and Ror2 was quantified in control and Ror1/2-KO fibroblasts after Control or TGF-β1 treatment.

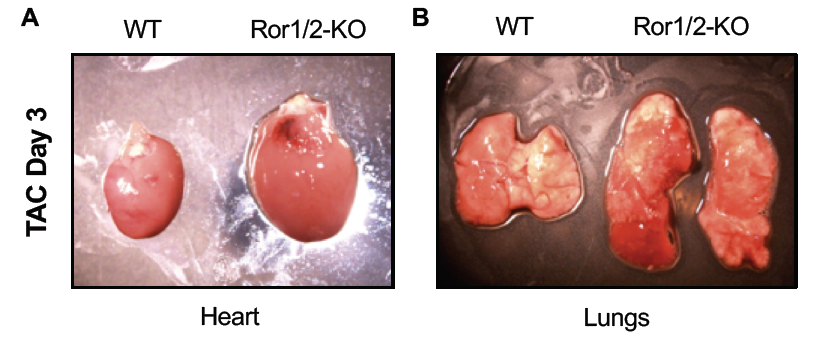

**Supplemental Figure 6.** Tissues from control and transgenic Ror1/2 double knockout mice 3 days post-TAC were isolated and imaged: A) Heart, and B) Lungs.
