## Supplemental Table 1 for "The Cell Surface Receptors Ror1/2 Control Cardiac Myofibroblast Differentiation"

**Table S1**. Primers for Q-PCR analysis of murine genes.

| **Gene** | **Forward Primer** | **Reverse Primer** | **Ref.** |
| --- | --- | --- | --- |
| Fsp1 | CTTCCTCTCTCTTGGTCTGGTC | TTTGTGGAAGGTGGACACAA | ^80^ |
| Acta2 | ACTCTCTTCCAGCCATCTTTCA | ATAGGTGGTTTCGTGGATGC | ^80^ |
| Postn | AAGCTGCGGCAAGACAAG | TCAAATCTGCAGCTTCAAGG | ^80^ |
| Ror1 | AGTTCCTCATCATGCGATCC | CCTTGTGCACGAA­­­­GAAGTGA | n.a. |
| Ror2 | CAGACGGCAATCCTGCACT | GATGACCCTTCGTGGCTCTT | n.a. |
| Slug | CATTGCCTTGTGTCTGCAAG | CAGTGAGGGCAAGAGAAAGG | ^80^ |
| Snail | CTTGTGTCTGCACGACCTGT | AGGAGAATGGCTTCTCACCA | ^80^ |
| Ptk7 | TGGCACCTCAGGATGTTGTT | GGACGGCTTCGGTTAGTGAT | n.a. |
| Prickle1 | TGCTCAGGAGATCCAAGTCC | CTCTCTTCAAAGTGATACGC | ^81^ |
| Vangl2 | TGCTCATGGTGCTTGTCTTC | GGAGCTCCAGCAGAACTACG | ^82^ |
| Il6 | GCTACCAAACTGGATATAATCAGGA | CCAGGTAGCTATGGTACTCCAGAA | ^21^ |
| Tnfa | CGGAGTCCGGGCAGG | GCTGGGTAGAGAATGGATGAA | ^21^ |
| Il1b | TGACAGTGATGAGAATGACCTGTTC | TTGGAAGCAGCCCTTCATCT | ^21^ |
| Ccl2 | CAGCCAGATGCAGTTAACGC | GCCTACTCATTGGGATCATCTTG | ^21^ |
| CD31 | GAGCCCAATCACGTTTCAGTTT | TCCTTCCTGCTTCTTGCTAGCT | ^83^ |
| CD45 | GGGTTGTTCTGTGCCTTGTT | CTGGACGGACACAGTTAGCA | ^83^ |
| Actb | AGAGGGAAATCGTGCGTGAC | CAATAGTGATGACCTGGCCGT | ^83^ |
